# A bacterial dynamin-like protein confers a novel phage resistance strategy on the population level in *Bacillus subtilis*

**DOI:** 10.1101/2021.07.26.453760

**Authors:** Lijun Guo, Laura Sattler, Peter Graumann, Marc Bramkamp

## Abstract

*Bacillus subtilis* DynA is a member of the dynamin superfamily, involved in membrane remodeling processes. DynA was shown to catalyze full membrane fusion and it plays a role in membrane surveillance against antibiotics. We show here that DynA also provides a novel resistance mechanism against phage infection. Cells lacking DynA are efficiently lysed after phage infection and virus replication. DynA does not prevent phage infection and replication in individual cells, but significantly delays host cell lysis, thereby slowing down the release of phage progeny from the host cells. During the process, DynA forms large, almost immobile clusters on the cell membrane that seem to support membrane integrity. Single molecule tracking revealed a shift of freely diffusive molecules within the cytosol towards extended, confined motion at the cell membrane following phage induction. Thus, the bacterial dynamins are the first anti-phage system reported to delay host cell lysis and the last line of defense of a multilayered antiviral defense. DynA is therefore providing protective effects on the population, but not on single cell level.

## Introduction

The dynamin-like protein (DLP) DynA in *Bacillus subtilis* is a member of the dynamin superfamily of proteins. DynA forms an unusual two-headed DLP that arose through a gene fusion in firmicute bacteria (1). Consequently, DynA contains two GTPase domains. The protein is involved in membrane remodeling processes and was shown to tether membranes in trans to promote full membrane fusion (1, 2). Deletion of the *dynA* allele in *B. subtilis* leads to a stress-sensitivity phenotype. We have recently shown that DynA plays a role in membrane surveillance (3). The protein is highly mobile at the membrane, but clusters into large and static assemblies upon addition of the pore-formation antibiotic nisin. Therefore, we hypothesized that DynA is recruited to sites of damaged membranes (3). Furthermore, a genetic link between DynA and the membrane organizing flotillin protein has been shown (4), supporting the notion that DynA is involved in membrane remodeling processes. We have previously reported that the *B. subtilis* phage □29 can form more plagues on the lawn of DynA-deficient strain compared with wild-type strain (3). □29 is the smallest phage known to infect *B. subtilis* so far, and its structure and the mechanism of its DNA replication have been extensively studied (5, 6). □29 forms tiny plagues on lawns of *B. subtilis* 168 even though it productively infects this bacterium in liquid medium (7). However, the molecular mechanism by which DynA interferes with phage infection remained unclear.

Recently, a variety of new bacterial resistance mechanisms against phage infection have been discovered. Phage infection imposes a tremendous pressure on bacteria to develop viral resistance strategies for survival (8–10). The evolution of bacterial phage defense systems in turn promotes the evolution of anti-resistance systems in phages and consequently, phages rapidly co-evolve to by-pass bacterial antiviral systems (11). Bacteria have evolved mechanistically diverse defense strategies that act at almost every stage of the phage infectious cycle. A first line of defense for bacteria is blocking phage attachment to the specific receptors, e.g. via production of extracellular matrix to occlude receptors, or via exploitation of competitive receptor inhibitors (12–19). Production of extracellular vesicles has also been proposed to act as a decoy mechanism to prevent phage attachment to cells (20). Also, block of phage DNA injection can be achieved by superinfection exclusion proteins that are commonly phage or prophage encoded (21, 22). However, the most abundant strategy to prevent phage infection is the specific degradation of viral nucleic acids by bacterial restriction modification systems, or by CRISPR/Cas systems that enable protection from invading DNA (23–27). While these mechanisms are acting on a single cell level and protect the individual cells, there are also systems acting on the population level. One of these mechanisms is abortive infection, which inhibits phage assembly in the infected cells by host cell lysis before mature viral particles have been assembled, thereby efficiently protecting the distribution of phages in the bacterial population (28, 29). Abortive infection, and thus early sacrifice of the infected cells, is among the most effective protection mechanism against rapid spreading of virus infections.

Here, we describe a novel phage resistance mechanism that prevents efficient host cell lysis. *B. subtilis* cells expressing the bacterial DLP DynA lyse later and with lesser efficiency compared to cells lacking this bacterial DLP. Single cell infection and phage replication are not affected by DynA, thus DynA provides a protective effect only on the population level. Upon phage infection, DynA forms large, almost immobile clusters at the cell membrane, likely preventing membrane rupture. Our data reveal that bacterial DLPs act as the first reported bacterial anti-phage system that blocks phage release from infected cells. DLPs are widespread in bacterial genomes and their combined activity against pore forming antimicrobials and their protective effect against virus dispersal in bacterial populations make them important parts of the bacterial stress response and innate immunity systems.

## Results

### DynA provides protection against phage infection

We have shown before that *B. subtilis* cells lacking DynA are more sensitive to phage infection by □29 and SPβ (3). We now aimed to unravel at which state of infection this protective effect is exerted. Therefore, we used the lytic *B. subtilis* phage □29 for infection assays. In line with our earlier observations we found that infection of *B. subtilis* with □29 causes small plaques. Infection of a *dynA* knockout strain (strain FBB002), however, results in a significant increase in plaque size (Δ*dynA, P* = 1.652×10^-46^) (**Fig. 1A**). Furthermore, the number of plaques was increased about four times when a lawn of *dynA* cells was infected compared to that of wild-type *B. subtilis 168* (*P*= 0.0014) (**Fig. 1A**), indicating that the lack of DynA renders cells more susceptible to lysis upon phage infection. Importantly, a strain in which we overproduced a DynA (here a functional DynA-GFP fusion in a strain background where the original *dynA* gene was deleted, strain FBB018) from an ectopic locus under control of a xylose inducible promoter (DynA++) did not produce any plaques after infection (**Fig. 1A**), indicating that an increased DynA level leads to a higher resistance. The observable effect of DynA on plaque formation becomes even more obvious with increasing cultivation time. The difference in the amount of phage induced plaques increased up to 25 h of incubation between WT and Δ*dynA* cells (**Fig. S1A**). After 25 h incubation, resistant bacteria started to form visible colonies and populated the plates again in wild type and Δ*dynA* cells, indicating that these cells acquired a resistance (**Fig. S1A**). We also analyzed the lysis behavior induced by □29 in liquid cultures. Exponentially growing cultures were infected with a MOI (multiplicity of infection) of 1 and optical densities were measured. Lysis of DynA-deficient bacteria was significantly faster than that of the wild type cells (**Fig. 1B**). We also wanted to test the effect of DynA overexpression. Therefore, we used a strain (FBB018) that expresses an ectopic copy of DynA (a DynA-GFP fusion). Cells were precultured with 1% xylose to induce DynA production, but xylose was removed during the infection experiment (DynA+). These cells lysed much later compared to the wild type cells and the optical density did not lower as much (**Fig. 1B**), indicating that the DynA proteins provided a protective effect. In order to increase the DynA amount further we also performed the infection assay with 1% xylose present throughout the infection (DynA++). The continuous overexpression of DynA reduced host cell lysis after □29 infection drastically (**Fig. 1B**). We concluded from this experiment that DynA has a protective effect, but may not entirely prohibit phage infection of cells.

**Figure 1.**
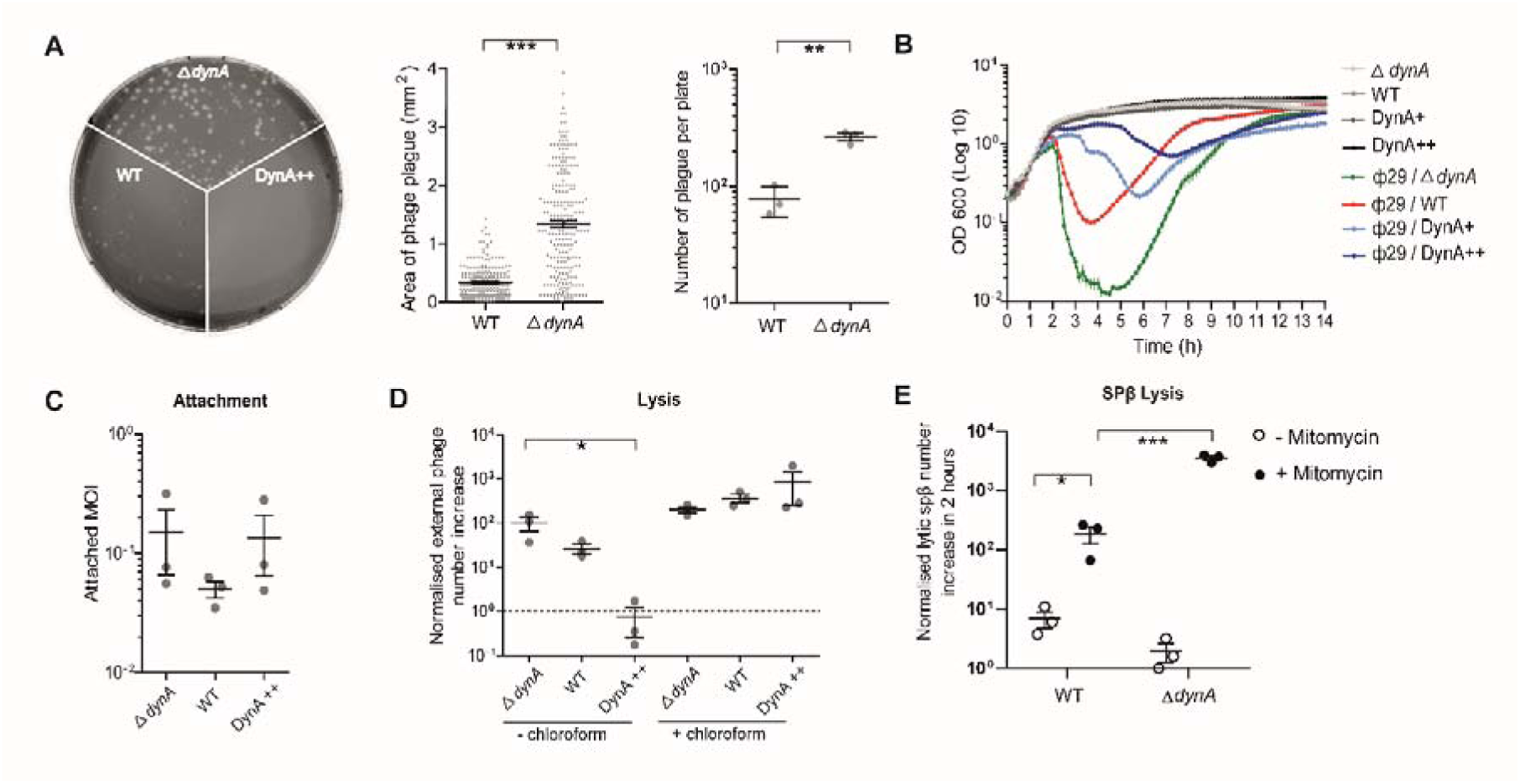
DynA confers resistance against phage infection. (**A**) □29 plaque formation on wild type *B. subtilis* 168 (WT), DynA-deficient strain (Δ*dynA*) and cells overexpressing DynA (DynA++). Note that almost no plaques can be observed when DynA is overproduced. Quantitative analysis of plaque sizes revealed significant larger plaques in Δ*dynA* compared to wild type. □29 plaque formation causes more plaques in Δ*dynA* cells compared to wild type. (**B**) The lytic effect of □29 infection (MOI = 1) can be suppressed by DynA overexpression. Optical densities of liquid growth cultures were measured in a plate reader. DynA+ was previously induced with 1% xylose for 30 min, but the xylose not present in growth experiment. For higher DynA expression levels 1% xylose was present throughout the infection experiment (DynA++). (**C**) Phage attachment in a mixture of bacteria and □29 (MOI = 1) was placed at 24°C for 10 min to allow phage attachment and then cells were separated from extracellular phages. Attachment rates were calculated based on quantitative spot assays. Note that there is no significant difference in phage attachment rates for wild type, Δ*dynA*, or DynA overexpression (DynA++) strains. (**D**) DynA overexpression reduces the number of released phages after infection. Phages were quantified by spot assays. Release of phages in the supernatant after infection (sample without chloroform) indicates a significant reduction of released phage progeny in cells overexpressing DynA. The total number of replicated phages (inside of host cells and released phages) was quantified after chloroform treatment. Note that no difference in total phage numbers in wild type, Δ*dynA*, or DynA overexpression (DynA++) strains is observed. (**E**). In cells lacking DynA (Δ*dynA*) the lytic induction of the lysogenic phage SPβ is significantly higher compared to wild type after 2 h mitomycin treatment. Statistical analysis is based on student t-test (* P value less than 0.05; two (**) less than 0.01; three (***) less than 0.001). Mean and standard error of three replicates are shown.

### DynA provides protective effects against phages and prophages

We next tested phage attachment and host cell lysis during the first infectious cycle for the three strains □wild type (168), Δ*dynA* (FBB002), and DynA++ (strain FBB018)] (**Fig. 1C**). A mixture of bacteria and □29 (MOI = 1) was placed at 24°C for 10 min to allow phage attachment. Subsequently, cells and attached phages were separated from free phages and the titer of free phages was tested by quantitative spot assays. Phage attachment was similar in all three strains (*P* = 0.3005 for Δ*dynA*: WT; *P* = 0.9129 for Δ*dynA* : DynA++). The average attachment rate of □29 was around 10 %. When the MOI was higher than 10, phages were attached to all cells (**Fig. S1B**). These results indicate that DynA does not significantly influence phage attachment. Next, we tested phage assembly and host cell lysis using quantitative spot assays (**Fig. 1D**). To this end, cells were mixed with phage □29 at a MOI of 1.0 and incubated at 37°C for 1 hour. Then, cells were harvested and the bacterial cell membrane was destroyed by chloroform treatment, thereby releasing the fully-assembled phage progeny. This method reliably lyses all bacterial cells independent of DynA concentrations, allowing us to compare a potential effect of DynA on intracellular phage assembly. After 1 hour of infection, the number of released phages and fully assembled phages was measured using quantitative spot assays. As a control we also determined the number of free phages that have been released independent of the chloroform treatment. In experiments without chloroform lysis of the host cells, the increase in active phages was highest for cells lacking DynA. However, the difference between Δ*dynA* and wild type was not significant for the three biological replicates we made. In contrast, in a strain overexpressing DynA (DynA++) we did not observe an increase in phage numbers during the infection and the difference in released phages between Δ*dynA* and an overexpression strain (DynA++) was significant (*P* = 0.0453 for Δ*dynA* : DynA++). When cells were lysed by chloroform correctly assembled phage particles that were not naturally released by host cell lysis, we observed no difference of phage numbers between wild type, Δ*dynA*, and DynA++ cells, indicating that DynA does not alter the □ 9 infection rate, or phage replication and assembly, but rather the release of mature phage particles.

The *B. subtilis* 168 strain we use for our experiments carries the SPβ prophage integrated into its genome. Upon cellular stress, such as DNA damage, SPβ is activated and assembled, leading to cell lysis (7). We were wondering whether the bacterial DLP DynA might only act against external, lytic phages or whether it may also have an effect on prophage proliferation. To address this idea, we performed a phage lysis test by inducing the lytic cycle of prophage SPβ by mitomycin C addition (**Fig. 1E**). The release of SPβ phage particles was tested by plating the supernatant from infection assays on a lawn of a SPβ-free *B. subtilis* strain 25152, carrying or lacking (PSB026) the *dynA* gene. Induction of SPβ in wild type 168 was observed and a low frequency of plaques was observed. However, when cells were treated with mitomycin C, the number of plaques increased significantly (*P*=0,0408 for WT:WT with mitomycin) (**Fig. 1E**). *B. subtilis* cells lacking *dynA* (strain FBB002) released a similarly small number of phages compared to the wild type when cells were not stressed by mitomycin C. However, when cells were treated with the DNA damaging agent mitomycin C, a large number of plaques were observed on a lawn of the susceptible LBJ17 cells (**Fig. 1E**). Therefore, absence of DynA resulted in a significant (*P*= 0.0005 for *dynA* with mitomycin:WT with mitomycin) increase of SPβ production and release. Thus, we conclude that the protective effect of DynA is not specifically act on □29, but seems to be a general mechanism interfering with viral release.

### DynA is not prohibiting phage replication in single cells

In order to precisely determine whether DynA only affects viral release and not phage replication or phage assembly, we used qPCR to quantify viral DNA in wild-type cells (168), Δ*dynA* (FBB002) and a DynA over-expression strain (FBB018) individually for internal, external and total phage DNA (**Fig. 2**). To test for the effectivity of phage infection we monitored again cell lysis in liquid cultures by measuring optical densities (**Fig. 2A**). When the MOI was equal to 10, the cell number of the Δ*dynA* strain dropped significantly faster compared to wild type around at 45 min post infection. In contrast a strain overexpressing DynA (FBB018) continued growing and showed no sign of massive cell lysis. Before 45 min of incubation, there was no significant difference in the growth rates between the three strains. We then analyzed the intracellular concentration of viral DNA (see material and methods) between the three strains. Within the first 45 minutes after infection the increase of viral DNA was similar in all three strains (*P* = 0.5189 for Δ*dynA* : DynA++; *P* = 0.9382 for Δ*dynA* : WT) indicating that DynA did not affect the replication of phage DNA. After 45 minutes the concentration of internal viral DNA was higher in the cells overexpressing DynA and lowest in cells lacking the bacterial DLP (**Fig. 2B**). To test whether the total phage production was similar in all strains, we quantified the total phage DNA and found that it remained indistinguishable within 90 min between all three strains (*P* = 0.9821 for Δ*dynA* : DynA++; *P* = 0.6218 for Δ*dynA* : WT)(**Fig. 2C**). This argues that phage replication (and hence viral DNA production) is not affected by DynA, but that cells lacking DynA release the viral DNA earlier into the medium, while cells with increased DynA concentrations seem to lyse slower and less efficient. If total phage production is similar, but cells lacking DynA lyse faster, we should find more viral DNA outside the cells in the medium. We therefore separated cells from medium and determined the concentration of viral DNA that has been released. In line with our hypothesis, we found more external viral DNA in Δ*dynA* cells and the lowest concentration in cells overexpressing DynA (**Fig. 2D**). Together, these data indicate that Δ*dynA* cells released the phage DNA approximately 45 min after infection more efficiently, while the lysis process of wild-type and over-expression strains were delayed. In summary, the presence of DynA did not influence phage attachment or viral DNA replication and phage assembly, but significantly slowed down host cell lysis and phage dispersal.

**Figure 2.**
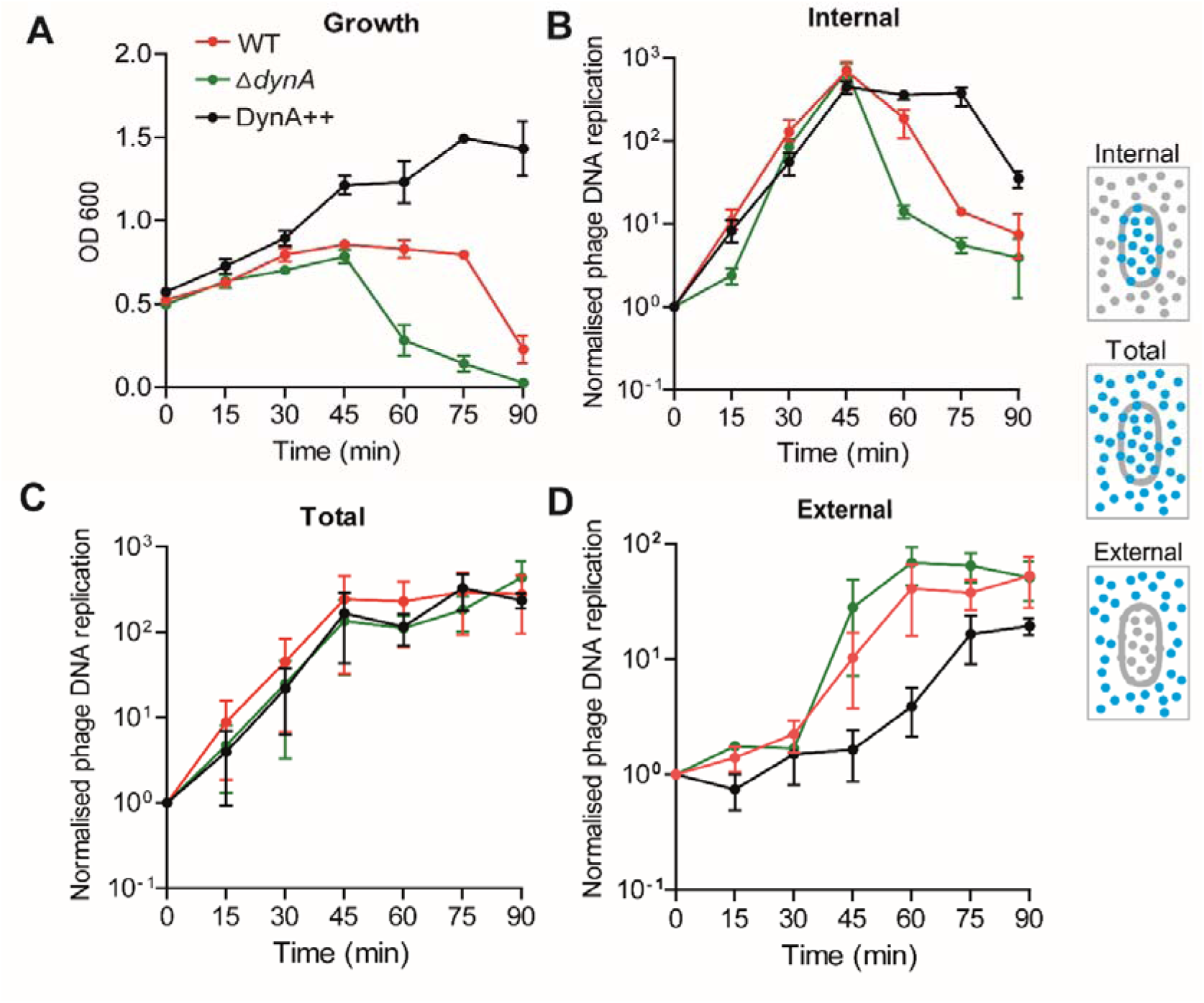
Phage DNA replication analyzed by qPCR. Phage DNA was quantified in *B. subtilis* wild type, Δ*dynA,* and the overexpression strain (DynA++) after □29 (MOI = 10) infection. The mixture of bacteria and □29 was pre-placed at 24°C for 10 min to allow phage attachment and then transferred to shaker of 37°C for measurement. Phage DNA of inside or on the cell, in total, and outside the cell were quantified every 15 min (see material and methods). (**A**) Optical density of infected cultures measured at 600 nm. Note the absence of a strong lysis effect in the DynA overproduction strain after phage infection. (**B**) Quantification of intracellular phage DNA. After 45 min the DynA overexpression strain contains more phage DNA trapped inside the host cell compared to the wild type and the Δ*dynA* strain. (**C**) The total phage DNA content in the samples is highly similar in all three experimental condition. (**D**) Quantification of external (released) phage DNA reveals that in cells overexpressing DynA less phages are released over the course of the experiment. Mean and standard error of three replicates are shown in all experiments.

### DynA does not colocalize with attached phages or viral DNA entry sites

We next wanted to visually follow the viral infection process and host cell lysis. Therefore, we fluorescently labeled phages and DynA *in vivo* to study the interaction between them on a single-cell level. To follow DynA dynamics inside the cell we used a DynA-GFP fusion protein expressed ectopically (from an *amyE* locus) under control of a xylose inducible promotor. The endogenous *dynA* gene in this strain was deleted (strain LGB03). As described before, in exponentially growing cells, DynA localizes around the cell periphery, associated with the plasma membrane. Addition of □29 phage lead to a change of DynA localization. Approximately 20 minutes after phage addition, but not before, DynA started to form clusters along the cell membrane (**Fig. 3A**). Thus, cluster formation did not occur during the early stages of phage infection, and hence, DynA clustering is not induced by phage attachment or DNA injection. We next labeled the phage DNA with the DNA-specific dye Hoechst. When we mixed phages with cells, we were able to visualize quickly cells that showed a small, bright blue fluorescent focus. During the course of infection, this focus disappeared and instead the entire cytoplasm of the infected cell was stained with Hoechst, indicating that the DNA dye is now dispersed throughout the infected cytoplasm. Since the bacterial nucleoid is degraded during virus replication (30), we assume that the majority of the stained DNA corresponds to viral DNA that is replicating in the host (**Fig. 3B**). The infected cell eventually lyses and the DNA bulges out of the host cell, which finally bursts and fragments, indicating dispersal of phage progeny (**Fig. 3B**). When we simultaneously followed DynA-GFP, and the cells were mixed with the DNA-labeled phage, we were able to observe that DynA aggregation occurred before host cells burst, releasing phage particles (**Fig. 3C**). We then wanted to label phage capsids and phage DNA simultaneously. To this end, we mixed CsCl-purified phages with Alexa Fluor 647 succinimidyl-ester (see material and methods). The succinimidyl-ester covalently attaches the dye to free amino groups of capsid lysine residues. These labelled phages retained their full infection potential and readily infected control cells. We used these capsid labelled phages for infection experiments using the strain expressing the DynA-GFP fusion protein (strain FBB018). Directly after mixing phages and cells, we observed a large number of phage particles close to the cell surface of the bacteria. However, DynA remained evenly distributed along the plasma membrane. During the course of the infection, the phage capsids gradually detached from the cells over time or decayed, leading to a disappearance of the far-red Alexa-647 signal (F**ig. 3D**). DynA clustering did not coincide with phage capsid localization during the early stages of infection, ruling out that DynA would react to phage attachment or viral genome injection. Finally, we wanted to track localization of viral capsids, viral DNA and DynA simultaneously. Therefore, we used the Alexa-647 labelled phages and stained their DNA with Hoechst. Also, the double labelled phages retained their full infection potential. When we infected DynA-GFP expressing cells, we saw that also at those sites where DNA was injected into the host cells (visualized by bright blue fluorescent foci), DynA did not accumulate. However, during later stages of the infection cycle DynA formed clusters that were enriched in the vicinity of phage egress sites (**Fig. 3E**). Taken together, these experiments argue for an involvement of DynA at sites in the cell membrane during release of the phage progeny. Missing colocalization of DynA with attaching phages and viral DNA injection, as well as the phage titer experiments, make it unlikely that DynA plays a role in prevention of infection at the single cell level.

**Figure 3.**
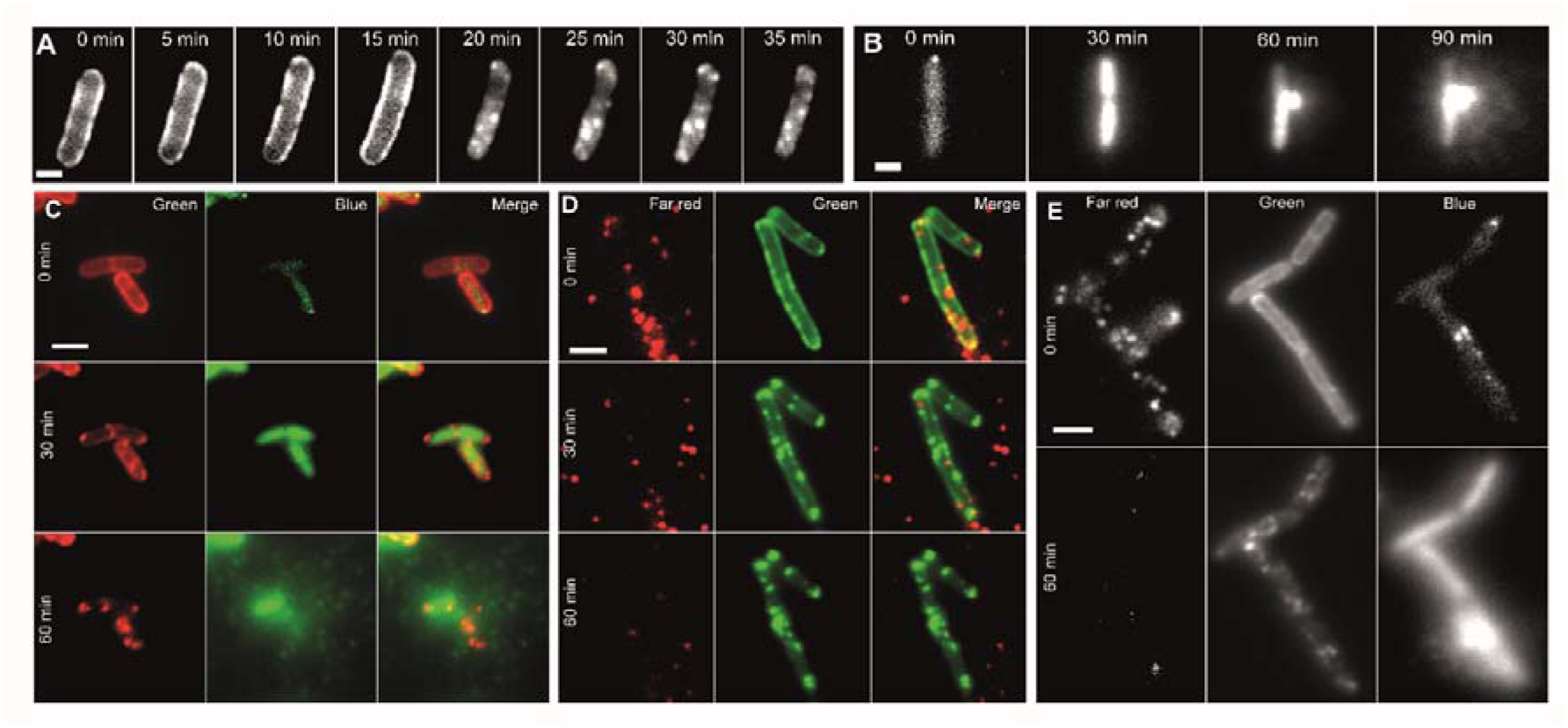
The in vivo dynamics of DynA during. c**29 infection observed by fluorescent labeling.** (**A**) □29 can induce DynA oligomerization *in vivo*. When the bacteria were exposed to phages (MOI= 1), DynA assembles into large clusters approximately 30 min after phage attachment at 37°C. Scale bar 1 µm. (**B**) Phage infection leads to host cell lysis. Phage DNA labelled with Hoechst DNA stain was used to track the infection progression. Upon attachment of the DNA-labelled phage (0 min) a small fluorescent focus can be observed on host cells. 30 min after infection the DNA stain has been injected into the host cell along the viral DNA. 60 min after infection cell lysis starts with bulging of cytoplasmic content including the fluorescently stained DNA. 90 min after infection the membrane bursts and DNA, likely including phage particles, is released. Scale bar 1 µm. (**C**) Analysis of DynA-GFP (false colored red) dynamics in infected cells. Phage infection is monitored by Hoechst dye injection into host cell by DNA-labelled phages (false colored green). Scale bar 2 µm. (**D**) Colocalization of □29 phages with labelled capsids (Alexa Fluor 647 dye, red) and DynA-GFP (green). Note that there is no correlation between localization of the phages and DynA at the onset of the infection cycle. Scale bar 2 µm. (**E**) Infection of cells expressing DynA-GFP (green channel) with double labelled phages [DNA stained with Hoechst (blue channel) and capsids labelled with Alexa Fluor 647 (far-red channel)]. Scale bar 2 µm.

### DynA delays and reduces host cell lysis

The results indicated that the protective effect of DynA on phage infection is unlikely happening at the stages of phage attachment and viral DNA replication. Thus, we reasoned that DynA may have an effect on host cell lysis and probably phage release. This would be in line with the biochemical activity of DynA in membrane protection by tethering and fusion reactions (1, 2). However, the extent of host cell lysis is difficult to quantify in different infection experiments, since there is a large variation in the degree of host cell lysis. To circumvent this problem, we designed an experiment in which cells deleted for *dynA* and cells expressing DynA-GFP were mixed. GFP fluorescence identified the cells that contain DynA, while cells without GFP fluorescence were Δ*dynA* cells. Fluorescence microscopy of the mixed cell cultures revealed that cells with and without DynA can be identified easily, while we did not observe fluorescence signals in the blue and far red channel (**Fig. 4A**). We than added the phages that we labelled on their capsids with Alexa-647 and the viral DNA with Hoechst dye. Immediately after mixing (0 min) we readily observed the far-red labelled virus capsids and, in few occasions, also small bright blue foci inside the *B. subtilis* cells (**Fig. 4A**, white square). A magnification of the cell shows that the Hoechst label is directly underneath an Alexa-647 signal, indicating that this is likely an event in which the phage DNA is injected into the host cell (**Fig. 4B**). The bulk of the bacterial nucleoid remained unstained at this stage of infection. After 30 minutes, most cells showed a dispersed Hoechst signal, indicating that by now most bacteria were infected and the Hoechst dye entered the host cells with the viral DNA. This allows an easy quantification of those cells that are infected. Together with the DynA-GFP signal we can quantify whether there is an infection bias between Δ*dynA* cells of DynA-GFP expressing cells. This was not the case, and thus, both strains were infected with the same rate. 60 minutes after infection we observed that the first cells lysed; indicative for lysis was the dispersal of the blue Hoechst dye diffusion. 90 minutes after infection many cells that lacked DynA (Δ*dynA*) were lysed, but most cells that contained DynA (DynA-GFP) were still intact and not lysed, yet (**Fig. 4A**). Finally, we compared the cell lysis ratio of Δ*dynA* cells and cells expressing DynA-GFP (**Fig. 4C**). We found that the ratio of cell lysis in Δ*dynA* was significantly higher compared to the cells expressing DynA-GFP measured 90 min post infection (**Fig. 4C)** (*P* = 0.0014 for Δ*dynA*: DynA++). These data clearly speak for a role of DynA in prevention of host cell lysis upon viral replication.

**Figure 4.**
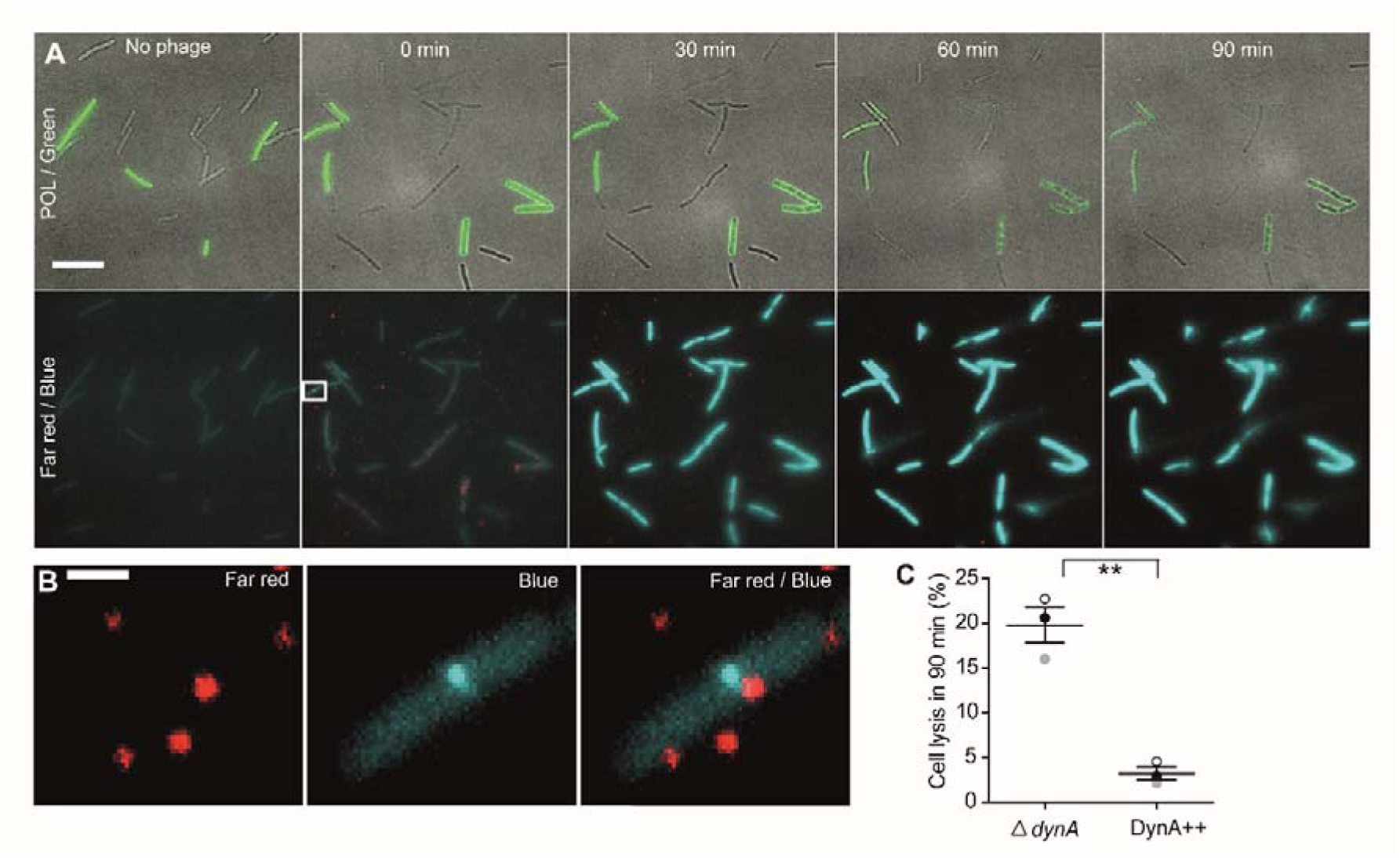
DynA prevents host cell lysis after infection. (**A**) Fluorescent microscopy analysis of a mixed culture of a DynA-overexpressing strain (DynA-GFP induced by 1% xylose, cells expressing DynA are indicated by green fluorescence) and a DynA-deficient (Δ*dynA*) *B. subtilis* strain after infection with □29 (phage DNA labelled with Hoechst dye and capsids labelled with Alexa Fluor 647). Time lapse analysis of the infection process reveals that shortly after addition of the labelled phages red foci appear next to the host cells and in few cases bright blue foci inside the host cells can be observed. Cells that are actively infected can be identified by the intracellular Hoechst DNA stain (blue). Cells with and without DynA are infected equally well. During the time course of the infection more of the cells lacking DynA lyse while cells expressing DynA remain intact. (**B**) Zoom into an early infection where the Hoechst dye injection into the host cell is spatially close to a labelled capsid, indicating an ongoing infection process. Note that no DynA focus is present at the site of DNA injection Scale bar 1 µm. (**C**) Quantification of bacterial lysis of DynA overexpressing (DynA++) and the deletion (Δ*dynA*) strains.

### Single molecule tracking analysis reveals DynA cluster formation upon phage infection

We also wished to bolster the findings of an increased formation of DynA foci at the cell membrane using single molecule tracking (SMT). In addition to statically positioned molecules visible by epifluorescence, SMT also visualizes and quantifies freely diffusing molecules. We used SMTracker software to analyze tracking data (31). If DynA was to diffuse throughout the cells, and would become more engaged in the repair of membrane irregularities in response to phage infection, we would expect a decrease of freely diffusing molecules and an increase in statically positioned molecules. We used 20 ms stream acquisition to track DynA-mVenus expressed from the original gene locus, under the control of the original promoter. A projection of all tracks (minimum length of four steps) was plotted into a standardized *Bacillus* cell of 3 x 1 µm size (**Fig. 5A**). While blue tracks indicate all freely diffusive molecules, red tracks show confined movement of molecules, and green tracks transitions between diffusive and confined movement. Clearly, DynA arrests at the cell membrane in some cases, while predominantly, it is freely diffusive. 60 minutes after infection with phages (MOI = 1), the number of confined, membrane localized tracks strongly increased, as expected (**Fig. 5B**). In order to further characterize the mode of diffusion of DynA, we employed Gaussian Mixture Modelling (GMM), in which displacements of molecules in x and y direction are evaluated as a probability density function. Tracks with little movement center around “0”, and fast tracks are away from the central axis. The shape of the function shown in **Fig. 5C** is clearly not Gaussian, which indicates the existence of at least two populations with different diffusion constants. In fact, data could be best described by assuming three distinct populations. These have diffusion constants of 0.023 µm^2^/s, 0.25 µm^2^/s, and 1.2 µm^2^/s, and sizes of 28, 45 and 26%, respectively (**Fig. 5E, F**). The populations are most easily explained by assuming a freely diffusive population, most likely consisting of monomers, a membrane-associated fraction diffusing in a constrained manner, and a slow moving/immobile fraction engaged in membrane repair. Upon phage infection, the static fraction increases to 38% of DynA molecules, to the expense of the freely diffusive molecules, while the intermediate fraction remained constant (**Fig. 5D, F**). This observation suggests that while more DynA molecules become actively involved in membrane repair, most molecules continue to scan the membrane for lesions.

**Figure 5.**
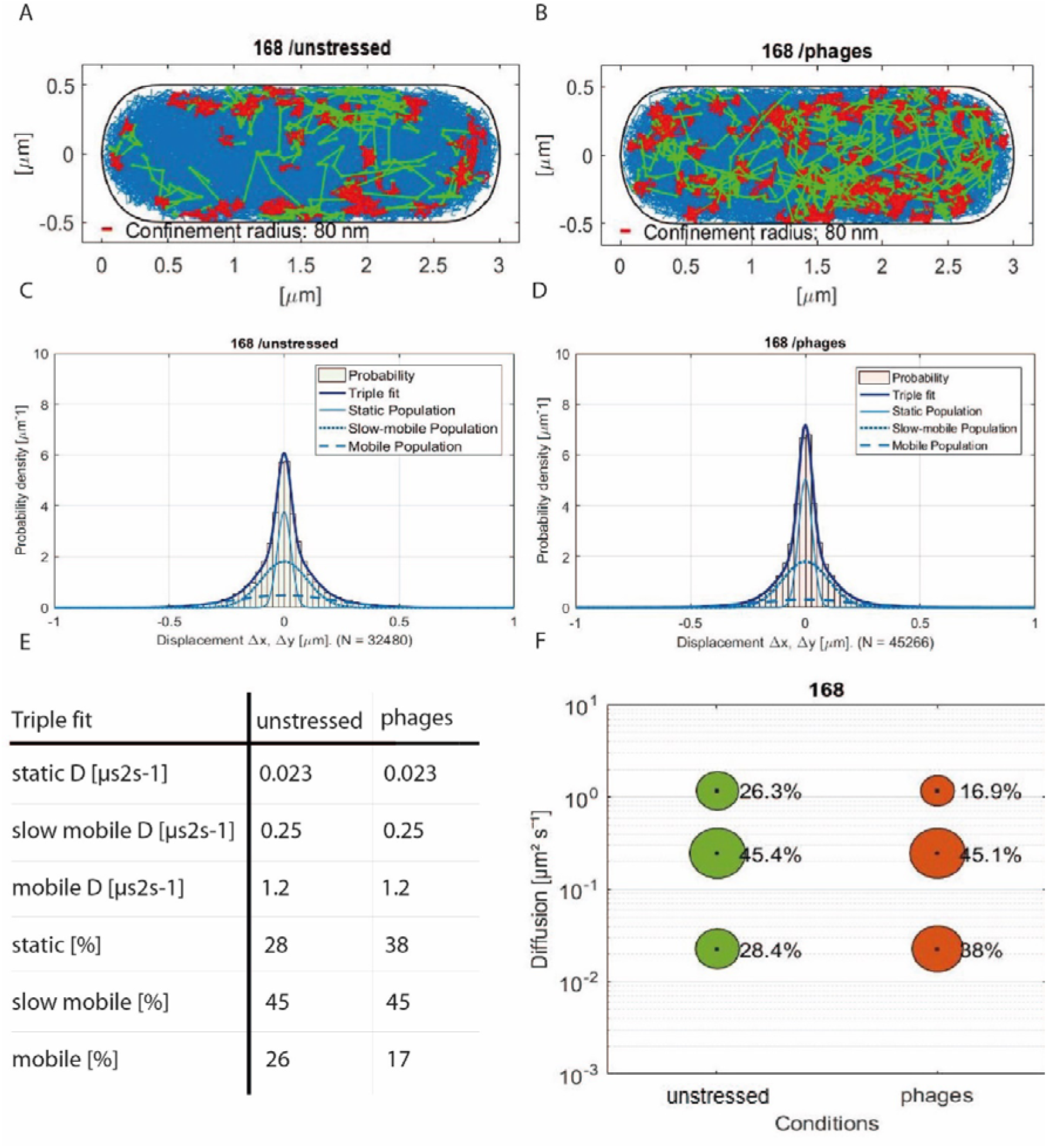
Changes of DynA dynamics in response to phage infection analyzed by single molecule tracking. Tracks of DynA-mVenus displayed in a standardized cell of 1 x 3 µm in (**A**) unstressed exponential growing cells or in (**B**) cells infected with □29 bacteriophages after 60 minutes. Freely diffusive tracks are shown in blue, confined tracks within a radius of 80 nm and a step length of 9 are depicted in red, and transient tracks showing mixed behavior are shown in green. (**C-F**) Diffusive behavior of DynA-mVenus in Gaussian-Mixture-Model (GMM) analyses of frame to frame displacements in *x*-and *y*-directions in exponential growth phase (**C**) 29 bacteriophages. The dark blue line indicates the overall fit of the three Gaussians distributions. Dashed, dotted and solid lines in brighter blue are the Gaussian distributions corresponding to mobile, slow mobile and static fractions, respectively. The diffusion constants ([µs^2^s^-1^] in (E) or *y*-axis in (F) and fraction sizes ([%] and bubble size) are shown in comparison to the two conditions of DynA-mV with a step size distribution of three populations, a static (lower bubbles), an intermediate mobile (middle bubbles) and a fast mobile (upper bubbles) fraction.

As a second measure for activity, we determined average dwell times of molecules, assuming that catalytically active molecules will remain in a confined motion, and thus statically positioned for many steps. Keeping in mind that dwell times are underestimated in our assays because of molecule bleaching during the acquisition, we can nevertheless conclude that following phage infection, dwell times of DynA strongly increase. During exponential growth, 87.7% molecules arrested for 73 ms (even freely diffusing molecules may stop for a short period of time), and 12.3% of the molecules arrested for 200 ms (**Table 1**); the latter will largely correspond to the molecules engaged in confined motion at the cell membrane (**Fig. 5A**). 60 minutes after infection, 71.3% of DynA molecules changed to longer dwell times of 290 ms, at the expense of molecules stopping for only 62 ms on average (**Fig. 5**). Thus, there is a strong shift towards extended dwell times 60 min after phage induction, supporting the idea that DynA molecules react to a strongly increased number of targets within the membrane.

**Table 1.** Average dwell times of DynA-mV

|  | **Exponential** | **Phages** |
| --- | --- | --- |
| average lifetime (frame/s) | 7.4 / 0.15 | 8.6 / 0.17 |
| average residence time<br>[s] | $0.100 \pm 0.011$ s | $0.283 \pm 0.011$ s |
| $\tau_1$ [s] | $0.073 \pm 0.00062$ s | $0.19 \pm 0.0062$ s |
| $\tau_1$ [%] | $87.7 \pm 1.32$ % | $28.7 \pm 5.23$ % |
| $\tau_2$ [s] | $0.2 \pm 0.012$ s | $0.29 \pm 0.0062$ s |
| $\tau_2$ [%] | $12.3 \pm 1.32$ % | $71.3 \pm 5.23$ % |

$\tau$ : dwell time

## Discussion

Bacteria have evolved a multitude of different phage defense systems to survive in a hostile environment where the number of phages exceeds the number of bacteria by an order of magnitude (32). Some of the most successful phage defense systems however, do not protect the individual infected cell, but rather act on a population level. This is the case for all abortive infection systems that lead to host cell lysis before the phage particles have successfully replicated (33, 34). We now describe a novel phage protection system that acts on the population level and exerts its function after phage replication and assembly has been completed. The presence of a bacterial dynamin-like protein, DynA in *B. subtilis*, interferes with the effective release of phage particles after infection. Thus, similar to abortive infection, dynamin-mediated delay of host cell lysis stops the phage epidemic from spreading rapidly through the population. Although abortive infection is the more powerful mechanism to reduce phage epidemics, the dynamin-mediated phage defense acts as a last line of defense to reduce the speed of the infection process in a bacterial population. Bacterial dynamins are therefore part of the multilayered bacterial response to phage infection with dedicated systems acting at each step of the phage infection cycle. Furthermore, the delay in phage particle release is independent of the phage species. We show here that DynA also exerts its protective effect against activated prophages such as SPβ.

Our data reveal that bacteria lacking DynA are infected at similar rates compared to their wild type siblings. Also, phage attachment and viral replication are unchanged in *dynA* mutant strains. Thus, lack of DynA does not result in an altered cell envelope structure that might reduce phage adsorption, thereby reducing infection rates. Rather, we show that presence of DynA at the cell membrane significantly reduces cell lysis and rupture of the plasma membrane. The protective effect of DynA becomes increasingly obvious, when this protein is overexpressed. It has recently been shown that phage lysis results in explosive cell rupture, but also a blebbing of the membrane with subsequent lysis has been observed (35). We frequently observed infected cells that had large bulges of cytoplasmic material (for example stained by DNA marker Hoechst). Large bulges of cytosolic content that is still surrounded by membrane suggests that these cells do not burst in an explosive and fast manner. We therefore hypothesize that DynA exerts a protective effect on the membrane that leads to a stabilization. This in turn reduces or at least delays the release of viral particles and explains the small plaque phenotype that we observe with cells expressing DynA. Overexpression of DynA enhances this effect drastically and cell lysis is largely reduced. We have shown before that DynA reduces phospholipid dynamics (3) and therefore there is likely a limit for the DynA concentration in the cell that still allows the required membrane fluidity homeostasis. Thus, there will be an tradeoff between fitness and a sufficiently strong protective effect against phage protection. Localization studies and single molecule tracking clearly show that DynA responds to phage infection. While DynA is highly mobile in uninfected cells, the percentage of freely diffusible DynA decrease and the percentage of stationary DynA at the cell membrane increases with infection. We have shown before that DynA is accumulating at sites of membrane damage (3). Thus, likely DynA is recruited to the membrane lesions caused by phage lysis and helps to maintain membrane integrity. We have shown before that DynA can tether and fuse membranes in a GTP-independent manner (1, 2). Scission and fusion of membranes is a common repair mechanism that is not only known for dynamin related proteins, but also seen with the ESCRT machinery (36). Similarly, it was recently described that the mycobacterial DynA homolog IniA is colocalized with sites of membrane lesion induced by cell envelope acting antibiotics (37).

We speculated before that GTP hydrolysis is required for the release of DynA complexes from the membrane (1–3). Since delay of host cell lysis by DynA does not lead to a survival of the infected cell, it seems logical that there is no selective pressure to recycle the DynA complexes after phage infection. Therefore, GTP hydrolysis is not essential for the protective effect that DynA exerts on phage infection. Nucleotide levels in dying cells are quickly dissipated and, hence, an active GTPase cycle would become impossible in lysing cells. The passive assembly and stabilization effect of DynA at the membrane, however, is very well in accord with the observed function in phage defense. The mechanism of action is therefore quite simple and does not involve sophisticated signal perception. A passive system that just reacts to membrane deformations is ideally suited to provide a last barrier against phage spreading. Also, a simple reduction of the explosive cell rupture will lead to a less effective phage distribution within the colony of bacteria, leading to a reduction of the epidemic spreading.

Dynamins are ubiquitous proteins and they seem to be early inventions in evolution. Stabilizing the plasma membrane against stress is a highly important feature and without dedicated membrane protection systems cellular life is unthinkable. In this aspect membrane protection can be compared to protection of genome integrity. It is therefore surprising how little we know about the molecular mechanisms that are used by cells to protect their cellular integrity. Dynamins seem to be ideally suited to fulfill this role and we should consider them not only as important proteins involved in membrane dynamics, but rather as a membrane protection component that contributes to the bacterial immune strategy.

## Acknowledgments

The authors acknowledge funding from the China Scholarship Council (CSC; fellowship to L.G.). Research in the groups of P.L.G. und M.B. was supported by a grant from the Deutsche Forschungsgemeinschaft (TRR174). M.B. acknowledges a grant from the Deutsche Forschungsgemeinschaft (BR2915/7-1).

## Author Contributions

L.G. performed all phage experiments and fluorescence microscopy. L.S. performed the single molecule tracking experiments. P.L.G conceived and evaluated the single molecule tracking experiments. M.B. conceived the study, analyzed the data and supervised the work. All authors wrote the manuscript.

## Declaration of Interests

The authors declare no conflicts of interest.

## Material and methods

### Strains and media

Bacterial strains used in this study are listed in **Table 2**. DynA-mVenus expressed from the original gene locus was constructed by cloning the last 500 bp (excluding the stop codon) of *dynA* into plasmid pSG1164 (38)containing the mVenus gene, using enzymes EcoRI and ApaI, after PCT amplification with forward primer GCTAGAATTCGACAACAGCCTTA and reverse primer GCATGGGCCCCATTTTTATTGTATTGTCTG. *B. subtilis* 168 was transformed with the resulting plasmid, establishing strain PG2112. The resulting Cells were grown in LB medium (10 g/l tryptone, 10 g/l NaCl, 5 g/l yeast extract) unless otherwise stated. Bacteriophages 29 and SPβ were purchased from the German Collection of Microorganisms (DSMZ GmbH).

**Table 2.** Strains used in this study.

| Strain | Genotype | Reference/Source |
| --- | --- | --- |
| *Bacillus subtilis* 168 | *trpC2* | Laboratory collection |
| *Bacillus subtilis* 25152 | *rpC2, metBI O, xin-1, SPβ-(erm)* | Laboratory collection |
| FBB002 | *trpC2, dynA::tet* | (1) |
| FBB018 | *amyE::Pxyl-dynA-gfp spc dynA::tet trpC2* | (1) |
| PSB026 | *trpC2, metBI O, xin-1, SPβ-(erm), dynA::tet* | (3) |
| PG2112 | *dynA-mVenus (cm)* | This study |

### Quantitative plague assay and spot assay

Overnight cultures of *B. subtilis* inoculated from glycerol stock were 100-fold diluted in fresh LB medium and grown to an OD_600_ of 0.5 - 1.0. Bacteriophages were diluted with 10-fold serial dilutions (1 to 10^10^) in LB medium. For plague assays, 100 µl of each phage dilution were mixed with 1 ml bacteria culture and the mixture spent 10 min at room temperature. Then with 4 ml LB with 0.5% agar were added to the mixtures and subsequently poured on LB agar plates. For spot assays, 1 ml bacteria were mixed with 4 ml LB complemented with 0.5% agar and poured first, then each phage diluteon (5 μl) was dropped to the plate. Plaques were detected after 6 hours of incubation at 37°C or overnight incubation (< 25 hours) at 24°C.

### Phage purification by isopycnic CsCl gradient centrifugation

1 liter of bacteria-phage mixture was prepared with a phage titer above 10^9^ PFU/ml. Bacteriophages were separated from bacteria by centrifugation at 3,800 g for 10 min. The supernatant contained bacteriophages and smaller cell debris. Phages were further purified by centrifugation at 13,000 g for 2 hours. Subsequently, 10 ml gelatin-free SM buffer (100 mM NaCl, 25mM Tris, 8 mM MgCl_2_, pH 7.5) was added to the pellet. The phage suspension was centrifuged again at 13000g for 10 min to separate cell debris. Solid CsCl was added slowly up to a final concentration of 1.40 g/ml, and swirled gently to dissolve in the phage suspension, The phage-CsCl solution was loaded to ultracentrifuge tubes and run 24 hours at 200,000 g using a Beckman 70.1 Ti rotor. The phage band was transferred to a 10 kDa cutoff dialysis cassette and dialyzed three times against 500 ml of gelatin-free SM buffer each. Finally, the phage preparation was filtered (0.45 μm) and stored at 4°C.

### Real-time PCR

Bacteria were grown at 37°C up to an OD_600_ 0.5 in LB medium. The culture was infected with □29 at a MOI of 10. All infected strains were incubated for 10 min at 24°C to allow phage attachment, then placed in 37°C shaker and timed. 500 μl aliquots of the *B. subtilis* cultures were withdrawn every 15 min. For analysis of the internal and external phage DNA concentration, cells and supernatants were separated by centrifugation at 3,800 g for 2 min and pellets were resuspended in 500 μl buffer. 50% chloroform (V/V) was added to interrupt the phage infection process and samples were centrifuged at 16,000 g for 10 min to remove cell debris. Analysis of the DNA samples were performed by real-time PCR in a Biorad 96-well real-time thermocycler (iCycler). For viral DNA amplification following oligonucleotides were used: □29-*gp8*-F: GTCAGGGCGATAACTTCA and □29-*gp8*-R: TACGATCAACAAGGGACG. The data obtained for each DNA sample was interpolated to a standard curve constructed with known amounts of purified, full-length □29 DNA. □29 DNA was isolated using Invitrogen PureLink genomic DNA Kit.

### 29 lysis test and lysogenic SPβ lysis test

In a □29 lysis test, *B. subtilis* strains were freshly grown to an OD_600_ of 0.5 in LB medium, then mixed with phage □29 at a MOI of 1.0 and incubated at 37°C for 1 hour. External (released) phages were collected by centrifugation at 3,800 g for 2 min. For the entire assembled phages (including the phages still inside the host cells), the samples were mixed with 1% chloroform (final concentration), then mixed by ten tube inversions, and finally centrifuged. A spot assay was performed on the lawn of a *dynA*-knockout *168* strain to measure the phage titers. For the SPβ lysis tests, 5 μg/ml mitomycin was added to the bacterial cultures of wild-type and *dynA*-knockout strains. Cultures were shaken at 37°C for 30 min. After incubation, mitomycin was washed away with fresh LB medium. Lysed SPβ were counted using a quantitative spot assay that was performed on the lawn of the SPβ carrying the *dynA::tet* allele (3).

### Phage-capsid staining

CsCl-purified phage (500 μl) were mixed with 5 μl Alexa Fluor 647 (Succinimidyl Ester, 1 mg/ml in DMSO) and rotated for 1 hour at 24°C. 1 M NaCl and 10 % PEG- 8000 were sequentially added to the solution and stirred for at least 30 min. The mixture was then centrifuged for 10 min at 13,000 g. The pellet was resuspended in gelatin-free SM buffer (> 1/5 of the original volume) and centrifuged again at 13,000 g for 10 min to obtain the supernatant. Labelled phages were separated from unbound dye using Illustra NAP-5 columns. Columns were equilibrated with 10 ml gelatin-free SM buffer and then load 500 μl of the supernatant. Every drop (∼ 50 μl) was collected in 200 μl PCR tubes and fluorescence intensity and phage activity was assayed using an Infinite200 PRO (Tecan, Grödig, Austria) fluorescent plate reader. Specially, 2 μl of drops were respectively added to 198 μl gelatin-free SM buffer in a 96-black plate and the fluorescence intensity [630/670 nm (Ex/Em)] Was measured. Another 2 μl of each fraction was added to 198 μl of a exponentially growing bacillus culture (OD_600_ = 0.5) in a 96-transparent plate and OD values were measured for 6 hours with 30-min intervals. Fractions containing active phages with high fluorescence based on the capsid labelling were collected and stored in dark at 4°C.

### Phage-DNA staining

CsCl-purified phages (500 μl) was mixed with 5 μl Hochest (1 mg/ml in H_2_O) and rotated for 1 hour at room temperature. 1M NaCl and 10% PEG-8000 were sequentially added to the solution and stirred for at least 30 min. The mixture was then centrifuged at 13,000 g for 10 min. The pellet was resuspended in gelatin-free SM buffer (> 1/5 of the original volume) and centrifuged again at 13,000 g for 10 min to obtain the supernatant. The NaCl/PEG clean-up was repeated twice.

### Fluorescent microscopy

*B. subtilis* strains were grown up to an OD_600_ of 0.5 - 1.0 in fresh LB medium at 37°C. Labeled □29 and DynA-GFP were visualized on a Delta Vision Elite (GE Healthcare) equipped with an Insight SSI illumination (Insight Lighting, Rio Rancho, NM, USA) and a CoolSnap HQ2 CCD camera (Teledyne Photometrics, Tucson, AZ, USA). Images were taken with an oil immersion PSF U-Plan S-Apo 1.4 NA objective. A 3- color standard set Insight SSI unit with excitation wavelengths (blue 390/18 nm, green 475/28 nm, far-red 632/22 nm), single bandpass emission wavelengths (blue 435/48 nm, green 573/36 nm, far-red 679/34 nm), and a suitable polychroic for blue/green/far-red were used. ImageJ (39) was used to analyze the micrographs.

### Single molecule tracking

*B. subtilis* 168 DynA-mVenus cells were grown in S7_50_ minimal medium at 30°C under shaking conditions to an OD of 1. Afterwards, the cells were infected with □29 bacteriophages for one hour, with a multiplicity of infection (MOI) of 1. Cells were spotted on coverslips (25mm, Marienfeld) and covered with an agarose pad (1% (w/v), made of S7_50_ Medium and a smaller coverslip (12mm, Marienfeld).

Imaging was performed with a Nikon Eclipse Ti microscope equipped with a high numerical aperture objective (CFI Apochromat TIRF 100XC Oil, NA 1.49), an EM-CCD camera (ImagEM X2, Hamamatsu) and a YFP filter set (BrightLine 500/24, Beamsplitter 520 and BrightLine 542/27). mVenus fluorophores were excited by the central part of a laser beam (TOPTICA Beam Smart, 515 nm, max. power 100 mW) with a laser intensity of 20 mW, corresponding to about 160 W/cm^2^. Each movie was recorded with an integration time of 20 ms via stream acquisition, using Nikon NIS-Elements BR. Movies consist of 3500 frames. Because single molecule level was reached after about 500 frames, the first 500 were discarded before analysis.

### Single molecule tracking – data analysis

First, the videos were cut with Fiji (ImageJ) (39) and the last 1000 frames were used. Afterwards, the cell meshes were set with oufti (40). For particle detection, U-track (41), a MATLAB software, was used. Here, the minimal length of tracks was set to 5 and to link to points, no gaps for the particle detection was allowed. With the MATLAB software SMTracker (31) the data was analyzed. Thereof the import panel - for localization and *dwell times* - and the Gaussian mixture model (GMM) analysis panel were used.

### Statistical analysis

We used the R suite program to calculate statistical significances (42) .

## Notes

### Competing Interest Statement

The authors have declared no competing interest.

