## Supplemental Material Figure S1 for "A bacterial dynamin-like protein confers a novel phage resistance strategy on the population level in *Bacillus subtilis*"

**for**

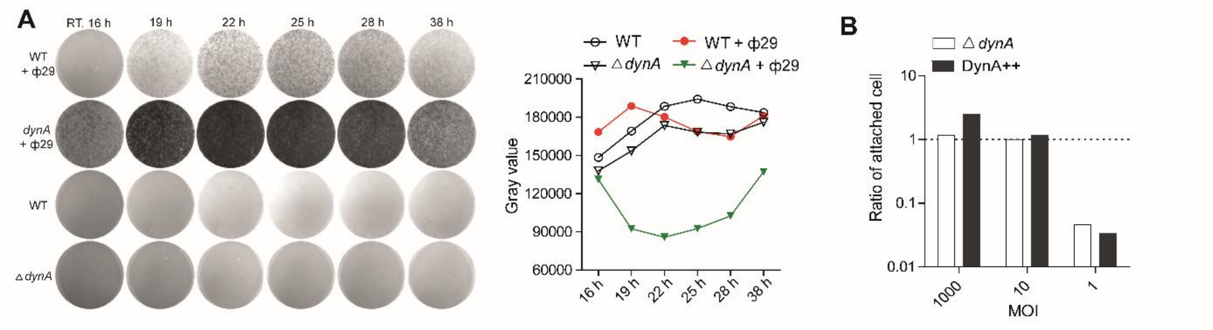


**Figure S1**. (**A**) Plaque formation of wild type and Δ*dynA* strains after infection with ϕ29 phage. Plates without phage infection are shown as control. Plates were incubated at room temperature and imaged for 38 h. Quantification of plaque formation was analyzed by gray values based on the pate images. (**B**) The proportion of bacteria attached to the phages after 10 min at 24℃ in experiments with varying amounts of phages (MOI 1000, 10 and 1).
